# Variations in the coverage of biological soil crusts along a gradient of aridity in the center-west of argentina

**DOI:** 10.1101/725986

**Authors:** A.L Navas Romero, M.A. Herrera Moratta, B. Vento, R.A. Rodriguez, E.E. Martínez Carretero

**Author notes:** These authors contributed equally to this work (A.L Navas Romero, M.A. Herrera Moratta). These authors contributed equally to this work (B. Vento, R.A. Rodriguez, E.E. Martínez Carretero).

## Abstract

The biological soil crusts (biocrust) play a fundamental role in the arid and semiarid areas of South America. However, little attention has been paid to the distribution and coverage of them. In Argentina, studies about biocrust are still scarce. The goal of this contribution is to analyze the coverage of the biocrust and each of the functional component along a gradient of aridity in the center-west of Argentina. The gradient included three differentiated sites: semiarid, arid, and hyperarid sites. The coverage was recorded using the Point-quadrat method on 30 transects through a gradient consisting of three sites: semiarid, arid, and hyper-arid sites. The arid site was the system with the highest coverage of biocrust followed by the hyper-arid site. The semiarid site had the lowest values of coverage and showed significant differences among the three systems were found. Cyanobacteria’s dominate in the hyper-arid site. On the other hand, cyanobacteria and lichens were dominant in the arid site. The coverage of studied organisms showed variations in the semiarid site. These results support the idea that the coverage has a strong relationship with the features of the studied ecosystem and the environmental factors both at a mesoscale and a microscale in a determined community.

## 1. Introduction

The arid, semiarid and hyper-arid zones occupy around 60 % of Argentina. Most of this extension corresponds to the Monte phytogeographic region (1,2). Biological soil crusts (biocrust) are fundamental components of the landscape, but they are scarcely studied in these ecosystems. The biological soil crusts are the result of an intimate association between microscopic poikilohydric organisms (algae, fungi-microfungi, and cyanobacteria) and macroscopic organisms (lichens, bryophytes, and micro-arthropods). These organisms are aggregated with soil particles, and they inhabit the first millimeters of the soil (3,4). The biocrust are widely distributed in many different kind of soils and in almost all plant communities where the sunlight reaches the soil surface (5), but they are dominant only in low productivity areas such as hyperarid, arid, semiarid, and polar sites (6).

Although the number of studies about biocrust have been increased, most of the researches on this topic become from the United States (5,7–10). Recently, many works about biocrust have been developed in Australia, China, Spain and Mexico (11–16). The biocrust play an important role in the arid and semiarid areas in South America. However, little attention was paid to the distribution of the biocrust, their taxonomic composition and even their coverage (17). In Argentina, studies about biocrust are still scarce and they have mainly focused on the impact of disturbances on the biocrust or the effect of the biocrust on the germination of vascular plants (18–20). Several biocrust communities have been described in the province of Buenos Aires (21), Chaco (22), San Luis (23) and the Patagonian region (19,24–28). Only a couple of recent studies in the Monte region have been made on grazing disturbances (18,29).

The increasing levels of disturbance in the ecosystems, the rate of disappearance of habitats, and the climate change threaten the persistence of the biocrust and make the collection of data about coverage essential. The biocrust are widely spread in the Monte phytogeographic province and play an essential role at the ecosystem level.

In this region, semiarid, arid and hyper-arid systems are located. In the latter, extreme environmental conditions result in areas of low vegetation coverage but allow the natural development of the biocrust (30,31).

We formulated the hypothesis the increase of temperature and decrease of rainfall from a semiarid to a hyper-arid system generate a change in the biocrust coverage. This may result in more development of the biocrust in the semiarid systems and a decrease in coverage when the aridity of the system increases. Together with changes in coverage, it is expected differences in the dominance of the functional groups from the biocrust. The results are the dominance of mosses in the semiarid system, and the increase of coverage of cyanobacteria in the hyper-arid system. Based on this, the goal of this contribution is to analyze the coverage of the biocrust and each of the functional components along a gradient of aridity in the center-west of Argentina. Understanding the differences in biocrust coverage along the aridity gradient contributes to elucidate what factors increase the probability of biocrust development. This knowledge allows the implementation of actions to protect this valuable ecological resource.

## 2. Materials and methods

### 2.1. Study site

During two consecutive years (2016-2017) which included two wet seasons (from December to February) and two dry seasons (from August to October), the biocrust coverage in a gradient of environmental stress in the Monte phytogeographic region located in the central-west of Argentina was surveyed. The gradient consisted of three sites: semiarid, arid, and hyper-arid. (Fig. 1).

**Figure 1.**
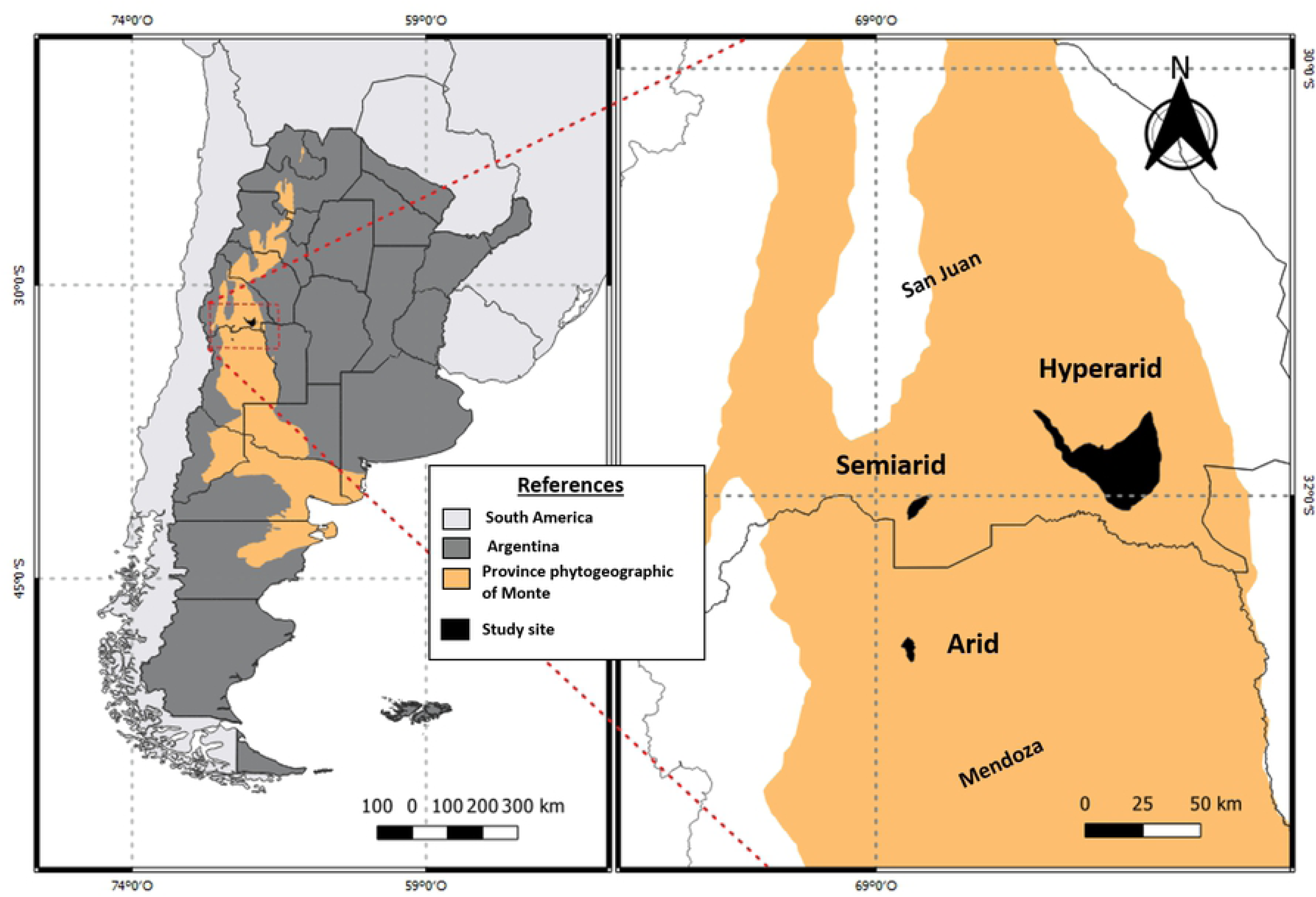
Map of the studied sites.

#### 2.1.1. Semiarid site

The semiarid site was located at the southwest part of the province of San Juan (32°00’8.43” S; 68°45’10.18” W) at 1139 m a.s.l. The site is part of the area called “Pedernal Protected Landscape” and it includes the conservation of 17,700 ha (32). The average annual rainfall is 370 mm and the average temperature 18 °C with a minimum of 6 °C, and a maximum of 20.7 °C (33). Soils correspond to Entisols in the typic Torriorthents (34,35). Dalmasso and Marquez (36) listed a floristic richness of 345 vascular species. The plant communities of *Zuccagnia punctata* Cav., *Larrea divaricata* Cav., *Baccharis salicifolia* Ruiz & Pav. together with *Hyalis argentea* Don ex Hook. & Arn. are dominant in this area.

#### 2.1.2. Arid site

The study area was located in the Capdevile district (32°43’24.3” S, 68°50’29.69” W), Mendoza province at 741 m a.s.l., in the Andean foothill. The average annual rainfall is 220 mm, 38 % of this rainfall occurs during the summer season (December-February). The average annual temperature is 17.5 °C, with a maximum of 30 °C, and a minimum of 3 °C (37). Soils correspond to Entisols in the typic Torrifluvents (38). According to studies made on the vegetation, *Larrea cuneifolia* is the dominant species in this area (39–41).

#### 2.1.3 Hyper-arid site

The study area was located in Los Médanos Grandes, in the province of San Juan (31°47’10.13” S, 67°58’55.75” W) on the eastern side of the Sierra Pie de Palo at 729 m a.s.l. The average annual rainfall is 103 mm and it occurs mainly from December (early summer) to May (autumn) (42). The average annual temperature is 18 °C with a maximum of 40 °C and a minimum of 10 °C (42). Soils correspond to Entisols in the typic Torripsamments (34). *Bulnesia retama* and *Larrea divaricata* are the dominant species, with a few representatives of *Prosopis flexuosa* DC. in depressed areas. Other accompaniment species are *Tricomaria usillo* Hook. & Arn., *Senna aphyla* (Cav.) H.S. Irwin & Borneby, *L. cuneifolia, Lycium* spp., *Atamisquea emarginata* Miers ex Hook. & Arn., and *Bougainvillea spinosa* (Cav.) Heimerl.

### 2.2. Experimental Design

The point-quadrat method (43) was used to determinate the presence and coverage of biocrust in each system and their functional groups (mosses, lichens, and cyanobacteria). A total of ten transects through an aridity gradient with surveys every 0.3 m were sampled. The first transect was randomly located, and the others were systematically located 10 m from the first one. This procedure was applied during the sampling of the four seasons.

Two periods (2016-2017) with their dry and wet seasons respectively were sampled and evaluated.

### 2.3. Data analysis

The statistical independence of the data was tested and the normality of the data was verified by the Kolmogorov-Smirnov test. The homoscedasticity was verified using the Levenne test. All analysis was performed considering α =0.05.

A multivariate variance analysis (MANOVA) and a Tuckey test of means were used to detect differences in the coverage of biocrust and among the main functional groups (mosses, lichens, and cyanobacteria) in the wet and dry seasons for each system. Additionally, differences among the systems were analyzed.

If data were no normally distributed a Kruskal-Wallis test was performed with a comparison of pairs. Graphics were made with the software “SigmaPlot v.11” (SigmaPlot, 2008). The statistical analysis was conducted using he program “Infostat v.16” (44).

## 3. Results

The three studied systems showed significant differences among them (H =61.92, p <0.0001). The arid system had the highest biocrust coverage, followed by the hyper-arid and the semiarid system, which presented the lowest coverage value. The cyanobacteria were the dominant functional group in the hyper-arid system. On the other hand, lichens were dominant on the arid system, while the semiarid system showed variations in the coverage percentage for the three analyzed groups. Significant differences were found among the three systems for lichens (H =60.31, p <0.0001). The hyper-arid, arid and semiarid systems showed significant differences among them, when considering the presence of cyanobacteria (H =22.35, p <0.0001) and mosses (H =24.48, p <0.0001).

### 3.1. Semiarid site

The semiarid system showed the highest value of coverage for vascular plants (V) 51.3 ± 11.8 %. However, the biocrust coverage value for this system was the lowest (21.1 ± 7.4 %) compared with the other two systems. The mean coverage value for the biocrust was higher during the dry seasons with (D): 23.7 ± 7.7 %, compared with the wet seasons (W) with 18.6 ± 5.9 %. During the 2016 period, the dry season (D1) showed the highest value of coverage 24.3 ± 7.6 % and the wet season (W1) the lowest one 18.23 ± 7.49 %.

Despite of these observed variations, no significant differences in the biocrust were found between the two seasons (Fig. 2A).

**Figure 2.**
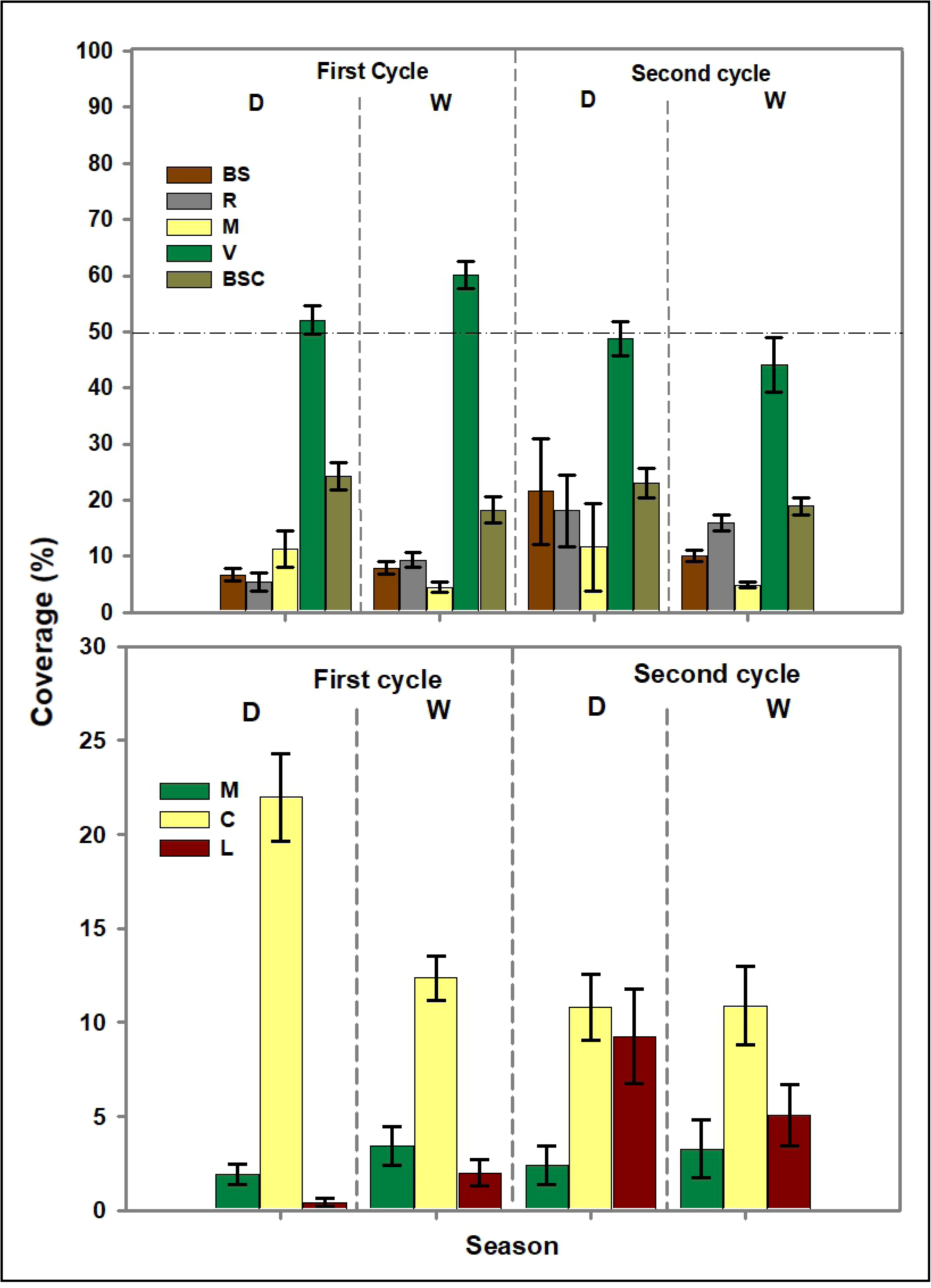
Mean percentage coverage for bare soil, rock, mulch, vegetation and biocrust (above) and mosses, cyanobacteria, and lichens (below) as a function of the seasons: D (dry season), H (wet season) in the semiarid system. The boxes represent the average values, and the bars represent the standard deviation. Different letters indicate significant differences between seasons (α =0.05). Bare soil (BS), rock (R), mulch (M), vegetation without biocrust (V), biological soil crust (biocrust), cyanobacteria (C), lichen (L), and moss (M).

The analysis of the dominance for each group resulted in a high mean coverage for the cyanobacterias with 13.2 ± 7.1 %, follow by lichens with 4.9 ± 5.7 % and mosses with 2.8 ± 3.4 %.

According to the analysis of the dominant organisms in each system, the group of cyanobacteria covers approximately 13.2 ± 7.1 % and this values was higher for all seasons compared with the coverage of lichens with 4.9 ± 5.7 % and mosses with a coverage of 2.8 ± 3.4 %. Significant differences were observed during the first dry season and wet season between cyanobacteria and lichens, and cyanobacteria and mosses. Moreover, significant differences were found during the second annual cycle between cyanobacteria and mosses, lichens and mosses, and cyanobacteria and mosses with the following values: W2 (H (D1) =17.01, p <0.0002, H (W1) =18.55, p <0.0001, H (D2) =10.26, p <0.0058, H (W2) =7.70 p <0.0211). On the other hand, lichens showed an increase in their coverage for the second annual cycle concerning the mosses.

In spite of the variations of the dominant groups in the biocrust, significant differences were found between seasons for lichens (D2 and W1, F = 3.32, p <0.0304) (Fig. 2B).

### 3.2. Arid site

The arid system showed the highest values of biocrust coverage (42.3 ± 14.2 %) from the three studied systems, even when compared with the coverage values of vascular vegetation (36.6 ± 11.9 %). The average coverage of biocrust was higher in the dry seasons with 43.3 ± 17.8 % respect to the wet seasons with 41.4 ± 14.5 %. However, no significant differences were found among seasons. The first dry season (D1) showed the highest values of coverage and the second one (D2) the lowest values. The temporary changes in coverage did not allow detecting significant differences between seasons (Fig. 3A).

**Figure 3.**
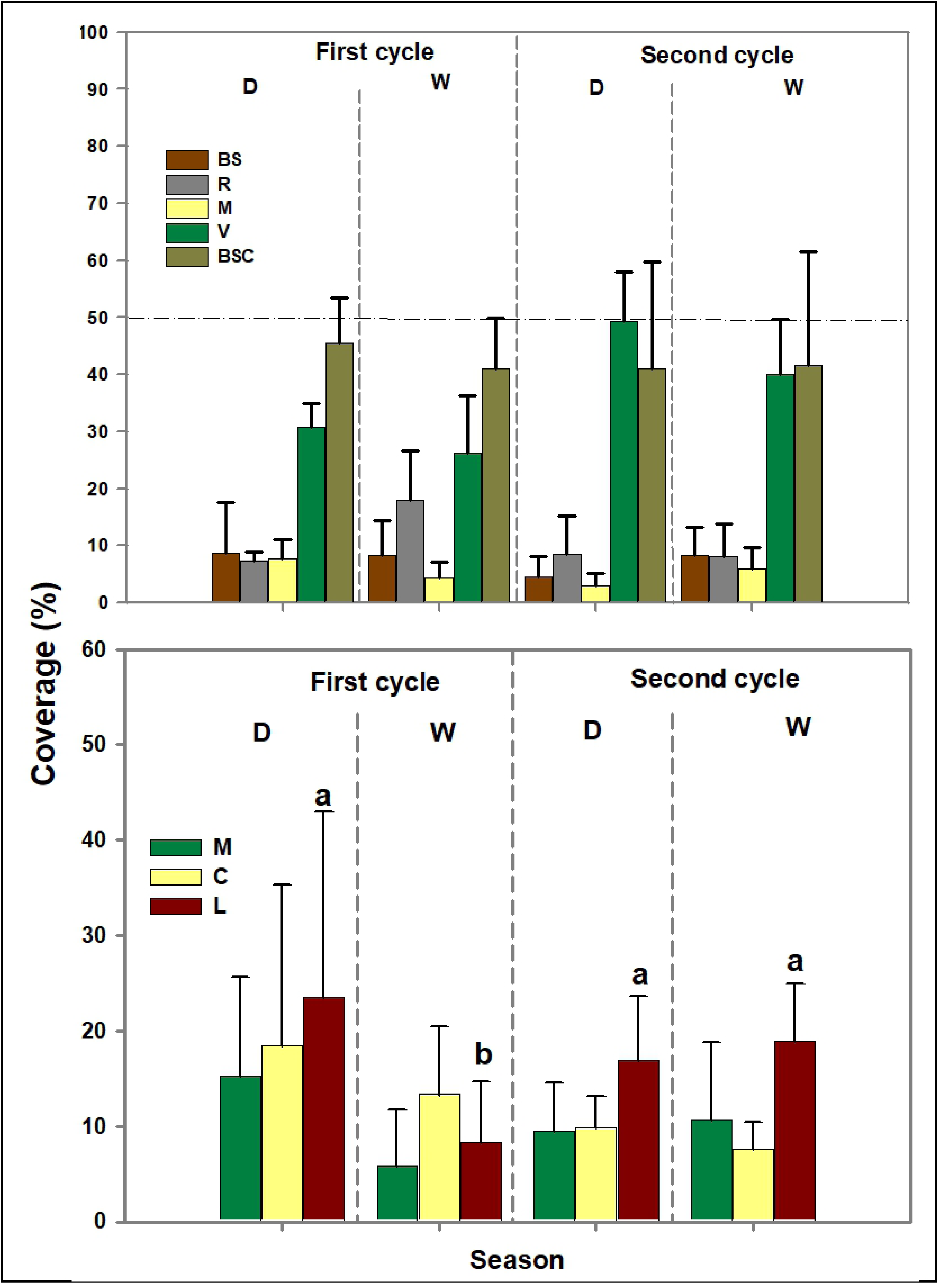
Mean percentage coverage for bare soil, rock, mulch, vegetation and biocrust (above) and mosses, lichens and cyanobacteria, as a function of the seasons: D (dry season), W (wet season) in the arid system. The boxes represent the average values, and the bars represent the standard deviation. Different letters indicate significant differences between seasons (α =0.05). Bare soil (BS), rock (R), mulch (M), vegetation without biocrust (V), biological soil crust (biocrust), cyanobacteria (C), lichen (L), and moss (M)

For the dominant groups, the coverage of lichens 16.9 ± 11.9 % was on average higher in all seasons respect to the coverage of cyanobacteria 12.3 ± 9.8 % and mosses 10.3 ± 7.9 %. For the wet season, the average coverage of the mosses was 8.25 ± 7.13 %, cyanobacteria 10.45 ± 5.91 %, and lichens 13.6 ± 7.9 %, while for the dry season for mosses was 12.4 ± 4.9 %, cyanobacteria 14.2 ± 3.3 %, and lichens 20.2 ± 6.4 %. Significant differences were found between groups for the second cycle, and only between lichens-cyanobacteria and lichens-mosses (H (D2) =6.19, p <0.0453, H (W2) =11.22, p <0.0036). Significant differences were found for the second year among groups. These differences were reflected between lichens-cyanobacteria and lichens-mosses group (H (D2) =6.19, p <0.0453, H (W2) =11.22, p <0.0036).

The cyanobacteria group showed a decrease in its coverage for the second annual cycle. The lichens and mosses did not show a clear and defined pattern of coverage. Even though the variations between dominant groups in the biocrust, significant differences were only found between seasons for the lichens (W1 with W2, D1, D2, F =2.96, p <0.045) (Fig. 3B).

### 3.3. Hyper-arid site

The hyper-arid system showed a coverage of vascular vegetation of approximately 50.4 ± 6.7 % and intermediate in the biocrust (25 ± 2.9 %) between the other two systems. The coverage of biocrust was higher in the dry seasons 27.5 ± 7.6 % in comparison with the wet seasons with a value of 22.5 ± 4.6 %. The first dry season (D1) showed the highest value of coverage: 27.3 ± 9.1 %, and the second wet season (W2) the lowest one (21.8 ± 5.3 %). Changes in the biocrust coverage showed seasonal variations and significant differences between the dry seasons (D1 and D2) and the wet one (W2) (H =8.39, p <0.03) (Fig. 4A). Respect to the dominant groups, the average coverage of cyanobacteria (19.6 ± 4.6 %) was higher in all seasons compared to the coverage of lichens (1.12 ± 1.32 %) and mosses (4.3 ± 2.4 %). Significant differences were found between cyanobacteria and mosses, cyanobacteria and lichens in D1 and W2. All the groups showed differences among them and seasons D2 and W2 (H (D1) =19.94, p <0.0001, H (W1) =23.42, p <0.0001, H (D2) =20.91, p <0.0001; H (W2) =20.2, p <0.0001). Additionally, all the groups showed an increase in their coverage for the second annual cycle. Significant differences were found between seasons for cyanobacteria (D1 with D2, W1, W2, H =10.48, p <0.01), lichens (W2 and D2 with W1 and D1, H =23.52, p <0.01), and mosses (D1 with D2, W1, W2, H =16.44, p <0.01) (Fig. 4.B).

**Figure 4.**
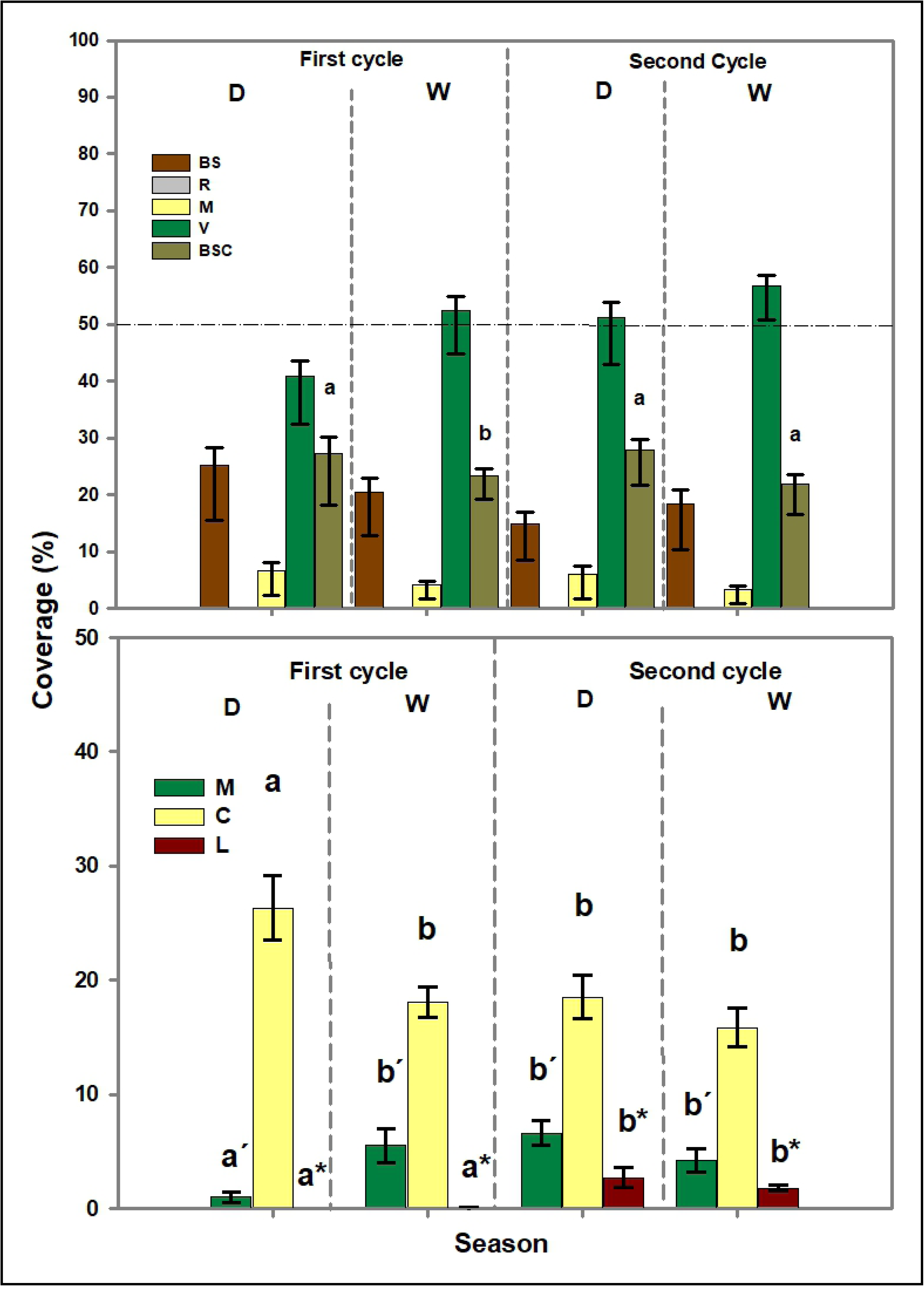
Mean percentage coverage for bare soil, rock, mulch, vegetation and biocrust (above) and for mosses, cyanobacteria, and lichen (below), as a function of the seasons: D (dry season), W (wet season) in the hyperarid system. The boxes represent the average values, and the bars represent the standard deviation. Different letters indicate significant differences between seasons (α =0.05). Symbols (‘, *) indicate different comparison levels. Bare soil (BS), rock (R), mulch (M), vegetation without biocrust (V), biological soil crust (biocrust), cyanobacteria (C), lichen (L) and moss (M).

## 4. Discussion

The biocrust were widely distributed in the three analyzed systems with the following coverage gradient: arid (A)> hyper-arid (H)> semiarid (S). The variations, at a microsite level in each system, such as soil texture, stoniness percentage, deposition and distribution of litter produce some limitations in the expansion of the biocrust (30). The coverage percentage of biocrust did not show significant differences among seasons which would indicate a relative stability of the communities. However, coverage was higher for dry seasons compared to wet seasons. Changes in biocrust coverage support the idea that the communities in the biocrust are more dynamic during short time periods (45,46). Results found in our contribution are coincident with those reported by Belnap et al. (2006), who observed a seasonal and yearly increase and decrease of the coverage of lichens and mosses when they evaluated the effect of temperature and precipitation on biocrust coverage.

The increase of biocrust during the dry season in desert areas could be related to the high temperatures that quickly would dry them. As a consequence of the dryness a deficit in carbon is produced until a net compensation point is reached (47). The summer season has high UV radiation and high soil temperatures, under these conditions soils become wet only for short periods which contribute to maintain or replace the damaged chlorophyll in organisms of the biocrust. In contrast, from late fall to early spring, there is a period of low UV radiation (little chlorophyll degradation) with low air temperatures that allow a longer up time in the biocrust and the chlorophyll can be synthesized. It has been shown that the biomass of cyanobacteria decreases during summer and increases in the late fall to early spring (48). Seasonal variations in biocrust coverage, with increases during the dry season, have been found in this work (Fig. 3) and by Belnap et al. (45,47).

Despite the fact of the detected changes, the biocrust turned out to be stable. The difficulty in detecting changes in the community on short time scales may be because some taxa in biocrust have a small size, are cryptic and have slow growth rates (46). These changes reflect some differences in the growth rate of each organism and its response to environmental factors. Cyanobacteria together with other unicellular components can colonize a site and rapidly expand the coverage in biocrust (49). Bryophyte coverage can increase faster in the absence of disturbance (50,51), and young mosses can grow more rapidly than old mosses (52). Lichens, generally, have slow growth rates that can decrease with age (53). Crustose lichens may invade shortly after disturbance (54) but may take a long time to increase their coverage. This statement suggests that a longer time period is necessary to identify changes in coverage and composition in communities of biocrust dominated by lichens and bryophytes.

The biocrust average coverage of the soil was 21 % for the semiarid, 25 % for the hyper-arid and above 40 % for the arid systems. In fact, data of the arid system, are coincident with that one’s reported by Belnap (49), who indicated that the biocrust reach to cover between 40-70 % of the surface in arid and semiarid zones at a global scale. The semiarid and hyper-arid systems (SH) presented a low coverage. However, these values are similar with that one’s reported by Castillo-Monroy and Benitez (55) for the biocrust of northern Mexico (21 %) and higher than the results reported by Molina et al. (56) for the northern zone of San Luis of Potosí, Mexico (9.37 %). The lower coverage found in the semiarid and hyper-arid could be related to the increase in soil stoniness and the reduction of the soil structure in these systems, which makes impossible the settlement biocrust, especially in inter-paths.

According to the recorded coverage of functional groups in the aridity gradient, it was found the following order: for cyanobacteria H > S > A, for mosses A > H > S, and for lichens A > S > H. These results are coincident with those reported by several works, suggesting that sandy soils as those present in the hyper-arid site are better for the development of cyanobacteria, while finer-textured soils (semiarid and arid sites) are more favorable for mosses and lichens (57–60). It is thought that the local settlement and distribution mechanisms of the bryophytes are related both to the capacity of the soil to retain water and to its higher stability in the face of disturbances (60). Anderson et al. (61) found that the abundance and diversity of bryophytes increased with soil clay content. On the other hand, Downing and Selkirk (62) made studies in the semiarid area of Australia and they found that the lowest bryophyte coverage occurred in sandy soils. In consequence, changes in the dominance of functional groups at a specific level, depends on multiple factors (15,30). The variation in the composition of the biocrust communities may also be related to the maturity of the biocrust, the distance to the vascular plants, the impact of temporal water availability, and increases in light and temperature at surface soil level, among others (5).

## 5. Conclusions

Communities of biocrust are widely distributed in the three studied arid systems. The coverage of biocrust is as follow: arid > hyper-arid> semiarid. The highest coverage of cyanobacteria was found in the hyper-arid system and the higher coverage of lichens and mosses in the arid system.

Although, on a macro-scale level, the environmental conditions exert influence on the organism type that dominates in a system; could be multiple factors (texture, stoniness, among others) that have influence on the specific composition of the CBS community on a micro-scale level. The differences in the composition and abundance of the different component groups may be due to the succession stage of the biocrust.

These results support the idea that the coverage, both at a macroscale and a microscale, depends on the characteristics of the studied ecosystem and the environmental factors in a determined community. This fact is relevant in the context of global change, because changes in the climate and the structure of the plant community can lead to critical variations in the functional structure and diversity of biocrust with serious consequences on the functions of the ecosystems. Changes in dominant groups could be reflected in the specific functioning of each type of biocrust in the system, affecting the ability of biocrust to keep water and protect the soil surface, influencing local erosion rates and hydrological and biogeochemical cycles in drylands.

## Acknowledgements

We thank Heber Merenda, José Vasquez, Yanina Rivas, and David Ponce by the assistance in fieldwork. This study was supported by the Consejo Nacional de Investigaciones Científica y Tecnológicas (CONICET).

